# Actinosomes: condensate-templated proteinaceous containers for engineering synthetic cells

**DOI:** 10.1101/2021.10.26.465899

**Authors:** Ketan A. Ganar, Liza Leijten, Siddharth Deshpande

**Affiliations:** Physical Chemistry and Soft Matter Chair Group, Wageningen University & Research, Stippeneng 4, 6708 WE Wageningen, The Netherlands

## Abstract

Engineering synthetic cells has a broad appeal, from understanding living cells to designing novel biomaterials for therapeutics, biosensing, and hybrid interfaces. A key prerequisite to creating synthetic cells is a functional three-dimensional container capable of orchestrating biochemical reactions. In this study, we present an easy and effective technique to make cell-sized porous containers crafted using the interactions between biomolecular condensates and actin cytoskeleton - we coin them actinosomes. This approach uses polypeptide/nucleoside triphosphate condensates and localizes actin monomers on their surface. By triggering actin polymerization at the expense of sequestered ATP and using osmotic gradients, the condensates are structurally transformed into containers with the boundary made up of actin filaments and polylysine polymers. We show that the GTP-to-ATP ratio is a crucial parameter for forming actinosomes: insufficient ATP prevents condensate dissolution while excess ATP leads to undesired crumpling. The surface of actinosomes lacks any structural order and is porous. We show the functionality of the actinosomes by using them as bioreactors capable of protein synthesis. Actinosomes are a handy addition to the synthetic cell platform, with appealing properties like ease-of-production, inherent encapsulation capacity, and an active surface which holds the potential to trigger signaling cascades and form multicellular assemblies, with potential for medical and biotechnological applications.

## Introduction

Cells are highly complex systems consisting of a plethora of interconnected biomolecular networks and this greatly limits our understanding of how they work. While deciphering molecular mechanisms in living systems is tedious, *in vitro* reconstitution assays is an excellent complementary approach to study specific cellular modules. In recent years, the bottom-up construction of “synthetic cells” has received tremendous attention, where compartmentalization is seen as an essential feature to mimic nature’s way of organizing reactions, and at the same time, providing a superior control^[1,2]^. Synthetic cells typically refer to an enclosed three-dimensional structure capable of performing tasks similar to their biological counterparts. Different types of synthetic cells have been proposed and can be broadly classified as membrane-bound and membraneless confinements^[3–5]^.

Membrane-bound compartments, built by the self-assembly of amphiphilic molecules have been widely used as cell-mimicking prototypes^[6]^. This has led to the design of a wide variety of confinements such as surfactant-stabilized water-in-oil droplets, liposomes with a lipid bilayer as the boundary, and even completely synthetic containers such as polymersomes and dendrimersomes^[5,7–9]^. These compartments are capable of reconstituting various biochemical processes within them and have been exploited to engineer a wide variety of cellular modules and to advance various applications like cell-free gene expression^[10,11]^, evolving proteins by directed evolution^[12]^, cytoskeleton assembly^[13,14]^, growth and division^[15–19]^, cargos for drug delivery^[20]^, and printing artificial tissues^[21,22]^. In these confinements formed via the hydrophobic effect^[23]^, membrane usually acts as a physical barrier and restricts passive transport of molecules across them. This is commonly resolved by incorporating transmembrane proteins like *α*-hemolysin, making them selectively permeable^[22,24,25]^. Additionally, newer strategies have been designed such proteinosomes which have a membrane comprised of cross-linked, amphiphilic protein-polymer conjugates^[26]^. Unlike the relatively inert membranes of liposomes and polymersomes, the proteinaceous boundary of proteinosomes can perform enzymatic reactions^[27]^. Methods to produce these confinements suffer various limitations: easy-to-use bulk bulk methods have poor process control and are prone to polydispersity (variation in the confinement size) and have a low encapsulation efficiency; employing microfluidic emulsion-based techniques effectively solve many of the issues, but at the cost of technologically advanced, sophisticated, and less accessible set ups^[28–30]^.

Biomolecular condensates, membraneless structures formed via the process of liquid-liquid phase separation have emerged as new types of bioreactors in recent years^[31]^. After their discovery and realization of the prominent role they play in intracellular biochemistry, they have been heavily exploited also in the realm of synthetic biology. Some salient features of condensates are their ability to sequester molecules and their assemblies^[24,32]^, resistant to extreme conditions^[33]^, support biochemical reactions with increased reaction rates and enhanced enzyme kinetics^[34–36]^ and exchange of molecules with their surroundings^[37]^. Interestingly, condensates have been explored as possible scaffolds to form synthetic containers^[38]^. For example, complex coacervates have been forged into multilayered compartments via a surface-templating procedure, albeit producing thick shells and the use of chemical treatments^[39]^. Another study demonstrated that the condensates formed by glutamic acid-rich leucine zipper and arginine-rich leucine zipper could be transitioned into hollow vesicles via temperature changes^[40]^. Alternatively, coacervate droplets can be coated with amphiphilic molecules; small unilamellar lipid vesicles were assembled at the interface of RNA/peptide droplets transforming them into an RNA encapsulated membrane-bound confinement^[41]^. These studies highlight the potential of condensates as templates to form novel confinements but also present several limitations such as thick shells, low membrane permeability, use of sophisticated protein engineering. If possible, one would desire a highly biocompatible proteinaceous confinement produced in a straightforward manner, without the use of complicated set ups.

In this study, we present a straightforward bottom-up approach to make cell-sized (2-6 μm) confinements with proteins as the building blocks. We start with condensates made up of a polypeptide (polylysine, polyK) and nucleoside triphosphates (NTPs), a mixture of adenosine triphosphate (ATP) and guanosine triphosphate (GTP). We then use actin, the well-known cytoskeletal protein capable of forming filaments, to structurally modify the condensate droplets. Actin localizes at the condensate interface and rapidly polymerizes into filaments at the expense high concentration of ATP present in condensates. Under the right conditions, this leads to internal coacervate dissolution, followed by complexation of polylysine with actin filaments at the surface, resulting in hollow containers which we term actinosomes. We show that ATP:NTP ratio is crucial in actinosome assembly and scanning electron microscopy reveals actinosomes as stable, porous containers. Finally, we show the capability of actinosomes as bioreactors by carrying out *in vitro* translation of proteins. We believe the addition of actinosomes, which can be formed without any use of sophisticated set ups and in a rapid manner, will be highly useful in the field of synthetic cells and to reconstitute reactions within cell-sized, biocompatible containers.

## Results

### Interaction of actin with multicomponent condensates forms actinosomes

We started with the idea of using membraneless condensates as a template to coat a biomaterial and subsequently dissolve the inner condensate to form a stable container (Fig. 1a). We aimed to bring about the structural and chemical transformation of the condensate by coupling a biochemical reaction, ideally carried out by the coated biomaterial itself. Complex coacervates made up of positively charged polypeptides (polylysine, polyK; polyarginine, polyR) and negatively charged NTPs (adenosine triphosphate) are widely used model systems^[42]^. With NTPs (ATP and GTP in particular) also being the common energy currency for a wide variety of biochemical reactions, we hypothesized that polyK/NTP would be a good starting point for our experiments. We determined the optimal concentrations of poly K and NTP to obtain maximum coacervation (Supplementary Figure 1). For all the experiments shown here, unless specified, polyK and total NTP concentrations were kept at 5 mg/mL and 5.4 mM, respectively. Using absorbance-based measurements, we estimated the amount of ATP inside the coacervates to be about 50 mM, *i.e.,* about 250 times more concentrated than the dilute phase, which was measured to be 0.19 ± 0.02 mM (see Methods for details). Our idea strengthened further when the addition of actin monomers to the system strongly partitioned them at the surface of these coacervates (Fig. 1b,c), similar to the observations made with other coacervate systems^[43]^. Based on fluorescence measurements, we calculated the partition coefficient of actin at the interface to be almost double (5.0 ± 0.9, *n* = 10) compared to its partitioning inside the coacervate (3.1 ± 0.29, *n* = 10). In a similar manner, the partition coefficient for polyK inside the coacervate was determined to be (3.7 ± 0.4, *n* = 10). In addition, we used a salt-deficient buffer, keeping the interfacial tension of the coacervate relatively high^[44]^. This also resulted in lower partitioning of actin inside the coacervate compared to the surface as actin readily partitioned inside the coacervates in presence of salt and did not accumulate at the interface (Supplementary Figure 2). Actin partitioning on the surface was also confirmed by measuring the surface potential of the coacervates through zeta potential measurements. We found the coacervates to be positively charged (16.9 ± 1.5 mV; Supplementary Figure 3), agreeing with previous observations^[45]^. We noted that the surface charge always remained positive irrespective of whether the polyK or ATP was in excess, suggesting accumulation of polyK molecules at the surface. Actin being net negatively charged at neutral pH was thus thought to assemble on the surface through electrostatic interactions ^[46]^. Indeed, surface charge measurements of actin-coated condensates showed significant reduction in the value of the zeta potential to 7.78 ± 1.13 mV within 10 minutes (Supplementary Figure 4).

**Figure 1.**
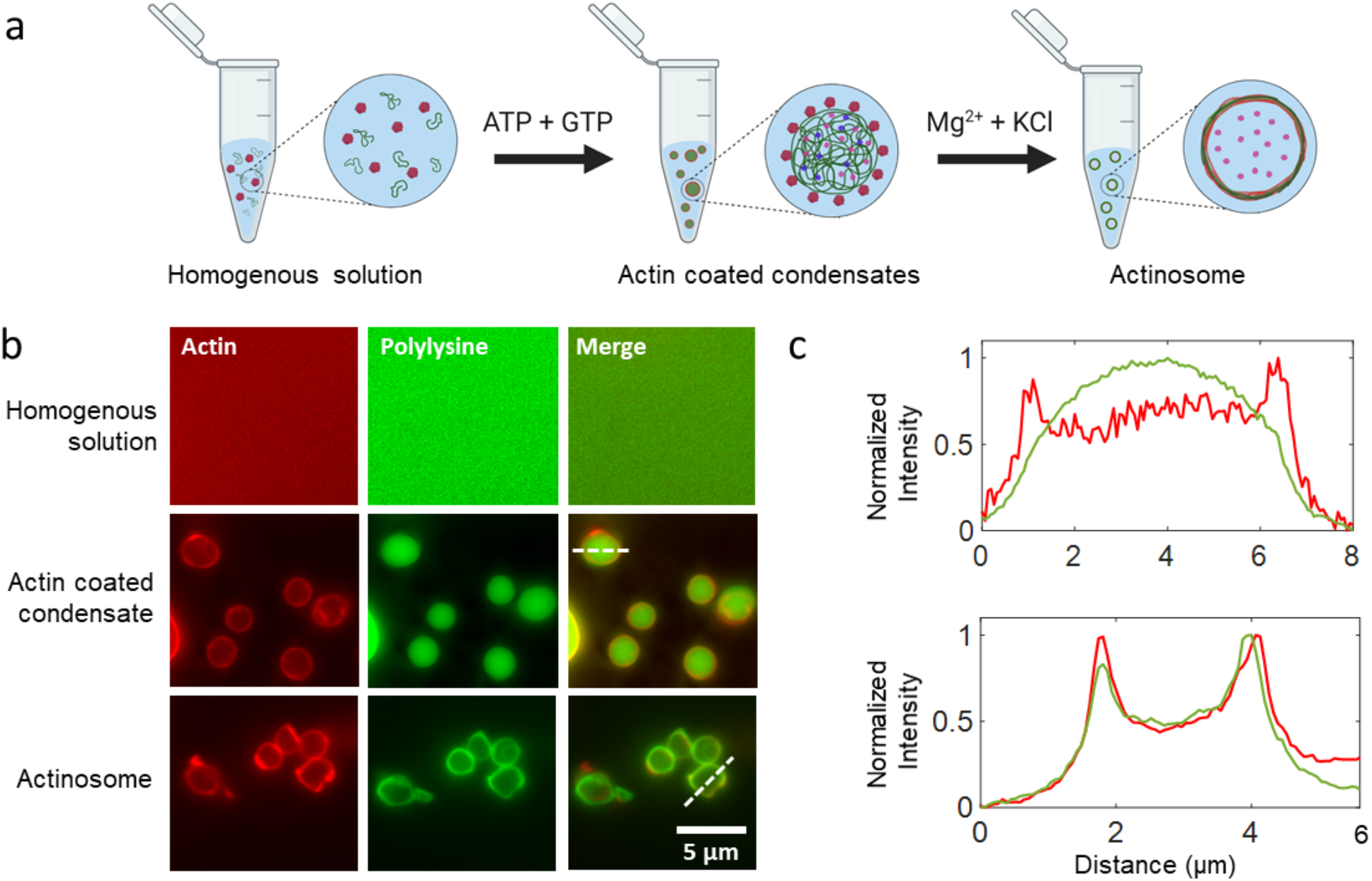
Condensate-templated actinosome formation. (**a**) A schematic demonstrating stepwise addition of reagents to produce actinosomes. **(b)** Top: A homogenous mixture of actin monomers and polylysine; middle: Addition of NTP mixture (GTP + ATP) triggers coacervation, resulting in polylysine/NTP coacervates with actin localized on the surface. Bottom: Mg^2+^ triggers actin polymerization at the expense of ATP hydrolysis, ultimately resulting in coacervate dissolution and formation of a shell made up of actin filaments and polylysine. **(c)** Line graphs (corresponding to the dotted lines in panel b) showing surface localization of actin on the condensates with polylysine concentrated in the interior (top) and colocalization of the filamentous actin and polylysine in actinosomes (bottom).

We triggered actin polymerization by adding a hypertonic buffer containing divalent cations (Mg^+2^) and KCl. This initiated the ATPase activity of actin, leading to a rapid hydrolysis of ATP present in the coacervates and formation of actin filaments on the condensate surface. To our pleasant surprise, when using an appropriate ratio of the ATP/GTP mixture, the condensates were subsequently converted into micron-sized somewhat spherical confinements within a matter of minutes. As can be seen from the fluorescence images in Fig. 1b, actin and polyK signals completely colocalized at the boundary of the (previously present) condensates, while the polyK signal from the lumen, indicating the presence of a phase separated environment, was significantly reduced. We aptly termed these containers actinosomes, where the actin filaments together with polyK polymers formed the container boundary, confining an aqueous lumen. The polyK localization also showed similar pattern of higher partitioning 2.9 ± 0.3 on the surface and 2.0 ± 0.2 inside the actinosomes. Based on the small increase in the dilute phase intensity, a finite fraction of polyK was assumed to leave the condensates altogether. As described earlier, the high salt content in actin polymerization buffer was necessary for actinosome formation. The hypertonic conditions likely decreased the interfacial tension, decreased the water content of the coacervates, and possibly facilitated outward movement of polyK filaments.

### ATP:NTP ratio is crucial to actinosome formation

Since ATP hydrolysis is crucial to coacervate dissolution and subsequent actinosome formation, we studied this further by tuning the ratio of NTPs. We maintained the total concentration of NTPs (GTP + ATP) constant at 5.4 mM and varied the amount of GTP from low to high, which we quantified as *R* = [GTP]/[NTP]. At *R* = 0, *i.e.,* when using only ATP, the coacervates immediately transitioned from a sphere to a collapsed state, resembling a crumpled structure, like a crumpled sheet of paper (Fig. 2a). This phenomenon can be explained as a combination of ATP hydrolysis and complexation of polyK with actin together with the osmolarity-induced water efflux leading to the buckling of the formed structure. We observed this crumpling prominently for *R* values below 0.6. Actinosomes were efficiently formed for R value between 0.7 and 0.8. As can be seen in Fig. 2b, the polylysine fluorescence rapidly decreased within a minute and co-localized at the interface along with actin. Thus, enough ATP was present for actin polymerization at the surface but at the same time, the inert GTP pool maintained enough osmolarity (~35 mOsm; hydrolyzed ATP possibly contributing further) preventing the crumpling and resulting in an actinosome with a wrinkled surface. The lack of a coacervate interior, judging by the lack of polyK fluorescence in the lumen but rather its colocalization with actin strongly suggests the presence of an aqueous lumen. Owing to the slight crumpling of the shell due to osmotic effects, the formed actinosomes were not perfectly spherical but were quite irregular in shape. We calculated the average size of actinosomes, by approximating them as ellipses, to be 3.1 ± 0.7 μm (major axis ± standard deviation; *n* = 41; Fig. 2c). We measured the eccentricity (major axis/minor axis) to quantify their spherical nature. A value of 1.2 ± 0.1 (*n* = 41) show that actinosomes remained reasonably spherical (Fig. 2d). At *R* values above 0.9, we observed a mixed population of both actinosomes and coacervates coated with actin. At *R* = 1, we observed only actin-coated condensates. With not enough ATP to bring about actin polymerization and coacervate dissolution, these coacervates remained stable and did not show any morphological changes over time. Thus, the ratio of GTP to ATP is crucial to actinosome formation.

**Figure 2.**
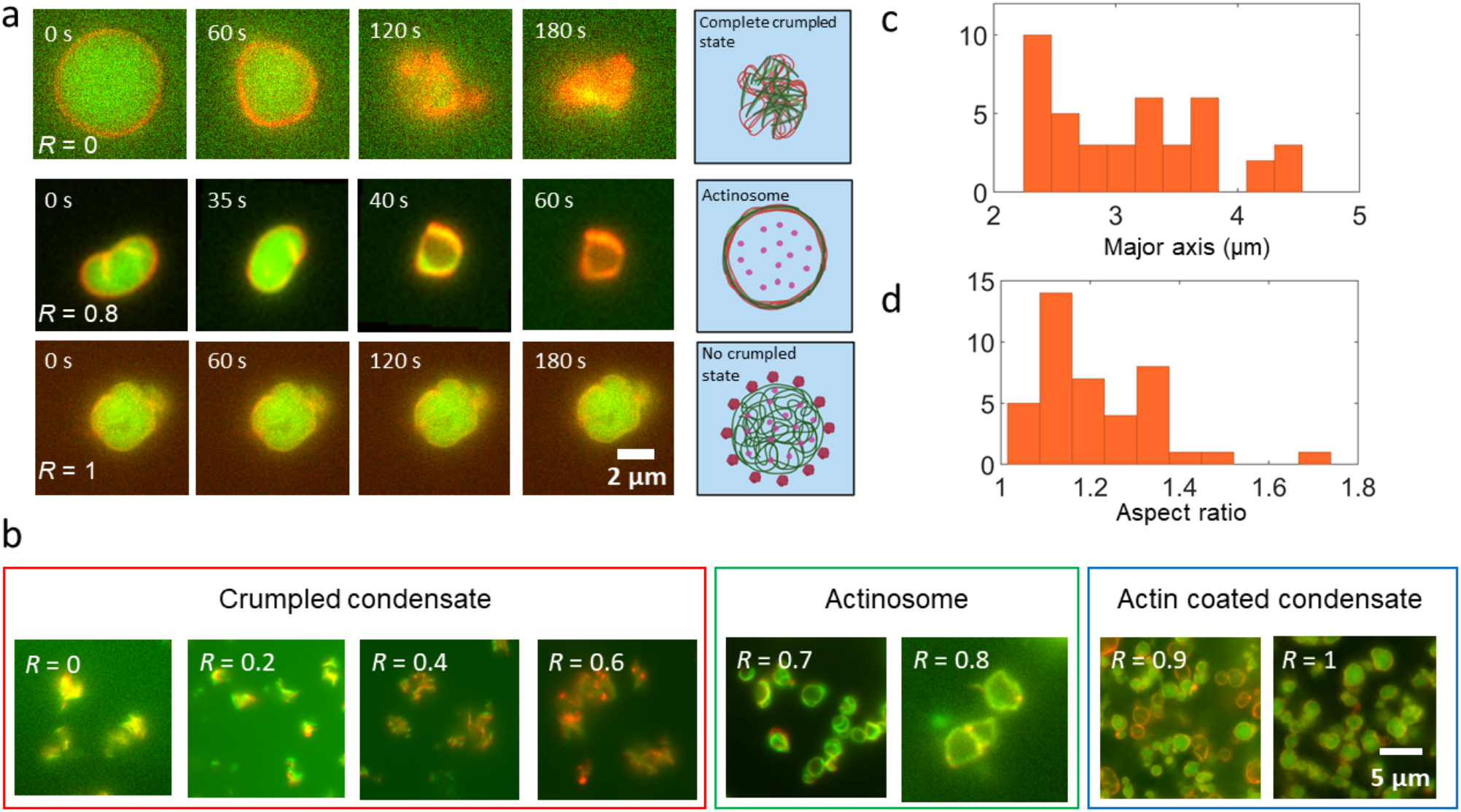
Actinosome formation depends on the ratio of NTPs present in the condensates. (**a**) Time-lapse images showing the actin-condensate dynamics at different *R* (= [GTP]/([GTP] + [ATP])) values. Low *R* values result in completely crumpled structures, intermediate values form cell-sized actinosomes, while higher *R* values result in stable actin-coated condensates. (**b**) Representative fluorescence images showing three key types of structures formed over the entire range of *R*. Actinosomes are obtained only within a narrow range (0.7 ≤ *R* ≤ 0.8). Lower values (*R* ≤ 0.6) result in crumpled structures while higher values (*R* ≥ 0.9) lack enough ATP and form stable actin-coated condensates. (**c**) Frequency histogram showing the size distribution of actinosomes, with mean size (major axis) of 3.1 ± 0.7 μm (n = 41). (**d**) Frequency histogram showing the ratio of major axis to minor axis; mean value of 1.2 ± 0.1 suggests that actinosomes tend to attain roughly spherical morphology (n = 41).

We also checked the effect of the nature of the polypeptide on actinosome formation, where we used poly-L-Arginine (polyR) to form the coacervates. While we obtained actin-coated condensates, we did not see complete crumpling when using only ATP neither a container formation when using a mix of ATP and GTP. This trend continued even after doubling the salt concentrations (Supplementary Figure 5). This might be due to significantly higher (100-fold) viscosity and surface tension (5.8-fold) of the polyR-containing droplets as compared to polyK-containing ones^[47]^, preventing rapid exchange of material across the interface and insufficient ATP diffusion to the surface.

### Actinosomes are hollow and porous containers

We then moved our attention to the surface characterization and the encapsulation efficiency of the formed containers. We performed scanning electron microscopy (SEM) studies to visualize the actinosome surface at high resolution. We prepared dried and sputtered samples of actinosomes (*R* = 0.74), crumpled condensates (*R* = 0.55), and actin-coated condensates (*R* = 0.92) for visualization (see Methods for details). The actinosome surface revealed a rough shell with no structural order of the filaments (Fig. 3a, b). The shells appeared thick, judging by the multiple layers that could be seen. They also appeared rigid, given that they survived the drying process and did not lead to any breakage. Importantly, small pores on the order of 0.2-0.5 μm in diameter were clearly visible, suggesting a permeable interface (Fig. 3b). On the other hand, the surface of polylysine/NTP coacervates (*R* = 0.92) with actin localized on the surface was smooth and did not show any of the above-mentioned features (Fig. 3c). At high ATP concentration (*R* = 0.55), crumpled structures were observed (Fig. 3d), corroborating with the fluorescence images obtained before. Given their porous nature, we wondered whether actinosomes could efficiently encapsulate charged biomolecules. We decided to encapsulate RNA, given its central importance in the cellular metabolism, and simply added it to the starting mixture of polyK and actin. We found that fluorescently (cy5)-labelled RNA (a 20-mer polyU) could be efficiently encapsulated inside the actinosomes (Fig. 3e). The partition coefficient of RNA was 4.0 ± 1.0 (*n* = 10) inside the actinosomes and much higher 7.0 ± 1.3 (*n* = 10) near the inner surface. Indeed, a line profile of the fluorescent intensity across the actinosome clearly showed the localization of RNA near the inner surface of the container (Fig. 3f). This is likely due to the electrostatic interaction between negatively charged RNA and positively charged polyK polymers, leading to a non-homogenous RNA distribution. We did not see any appreciable leakage of RNA fluorescence over time outside the actinosomes. Thus, despite their porous nature, actinosomes can efficiently encapsulate and retain biomolecules like RNA. We also checked the stability of actinosomes at lower temperatures, to gauge the possibility of long-term storage. We flash-froze the actinosomes in liquid nitrogen and stored at them −80°C. Upon reviving them at room temperature after 24 hours, their morphology looked similar to the ones that were freshly prepared (Supplementary Figure 6).

**Figure 3.**
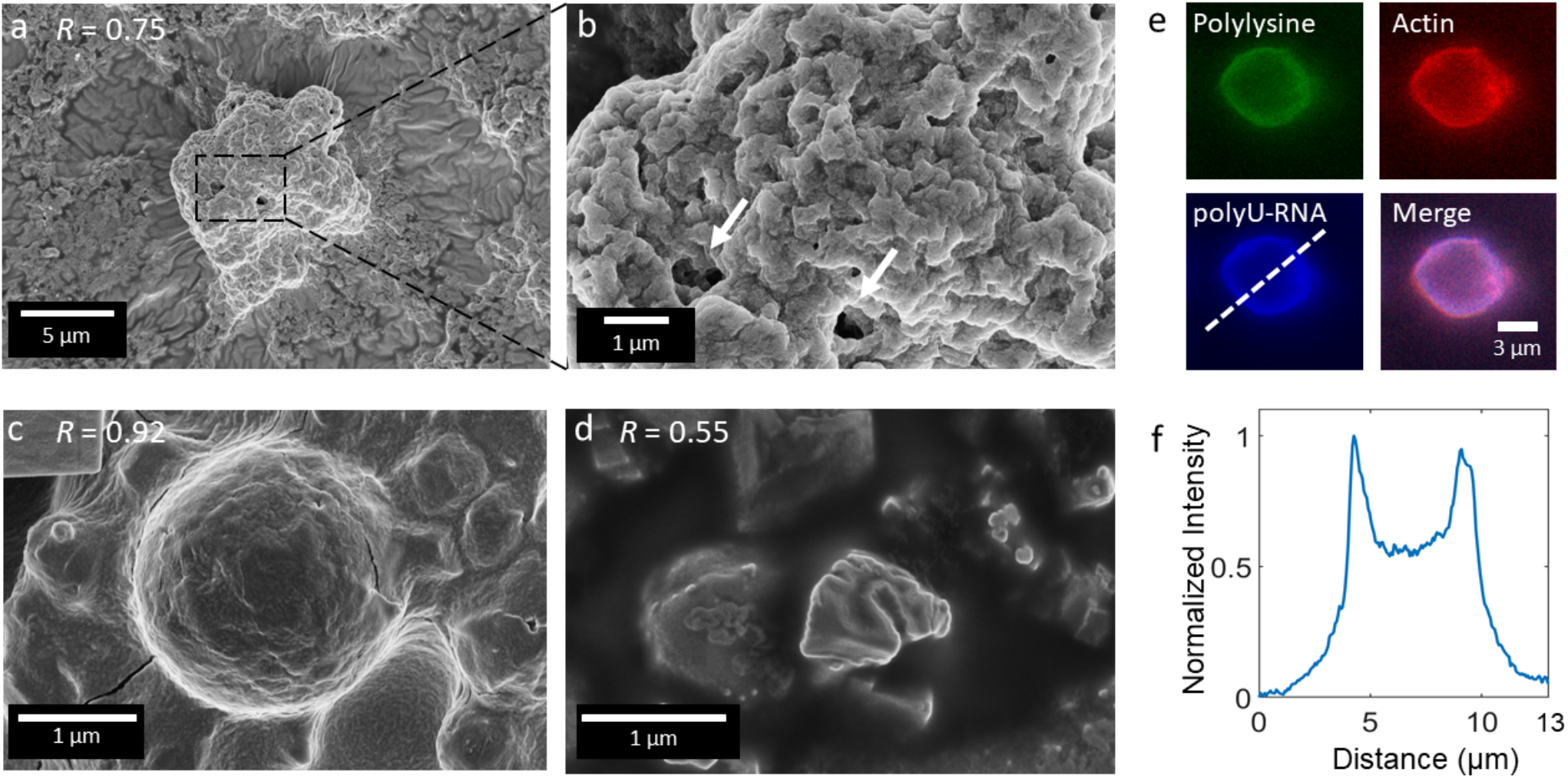
Actinosomes are porous containers and can efficiently encapsulate biomolecules. Scanning electron microscopy images of the structures formed at different *R* values. (**a**) Actinosomes (*R* = 0.75) appear as slightly crumpled spheres, similar to fluorescence images. (**b**) A zoom-in further reveals a rough, unstructured, multi-layered surface. Several sub-μm-sized pores are clearly visible (indicated by arrows). (c) Actin-coated condensates (*R* = 0.92) showing a spherical morphology and a smooth surface without any pores or structuration. (d) Crumpled actin-polylysine structure (*R* = 0.55) because of high ATP concentration present in the condensates. The obtained structure is comparatively smaller and shows several folded surfaces. (e) Encapsulation of Cy5-labelled RNA (polyU 20-mer) encapsulated inside the actinosomes. (f) Line graph(corresponding to the dotted lines in panel e) showing the localization of polyU-RNA at the inner surface of actinosomes.

### Actinosomes can efficiently carry out complex biochemical reactions

One of the trademark properties of coacervates is their ability to efficiently concentrate biomolecules within them, often up to orders of magnitude higher than the surroundings^[48]^. Coacervates also provide a distinct microenvironment that can differ from the dilute phase like concentration of metal ions such as Mg^2+[32]^. Thus, condensate droplets acting as the initial scaffolds for actinosomes provides an excellent opportunity to pre-load the actinosomes with components of interest. We tested this strategy by sequestering a cell-free protein translation machinery (retic lysate) along with single-stranded mRNA encoding green fluorescence protein (GFP) inside the coacervate droplets. This was simply done by adding the necessary components prior to condensate formation. Upon subsequent actinosome formation, we incubated the solution at 29°C and monitored GFP expression in real time (Fig. 4a). As can be seen, fluorescence in the GFP channel steadily increased over the course of an hour, with the protein expression evident as early as in the first few minutes. While fluorescence could be observed in the lumen, it was more intense at the shell, indicating the localization of the translation machinery or of the mRNA near the actinosome boundary. We did note some unwanted signal from the ATTO-532 labelled actin in the GFP channel, and an increase in the actin fluorescence itself because of the free actin in the dilute phase gradually accumulating on actinosomes. To confirm that GFP was indeed expressed, we conducted the same experiment with reduced fraction of the labelled actin (1 mol% instead of 10 mol%) and subsequently bleaching the fluorescent actin at the 25-minute interval by acutely increasing the light intensity for a short amount of time (Fig. 4b). This bleaching step completely eliminated the actin fluorescence but did not have any effect on the GFP fluorescent intensity confirming that GFP was expressed within actinosomes. Also, a negative control without the GFP-encoding mRNA, but with retic lysate, showed a faint signal in the GFP channel, coming from labelled actin, but it stayed constant over the entire duration (Supplementary Figure 7).

**Figure 4.**
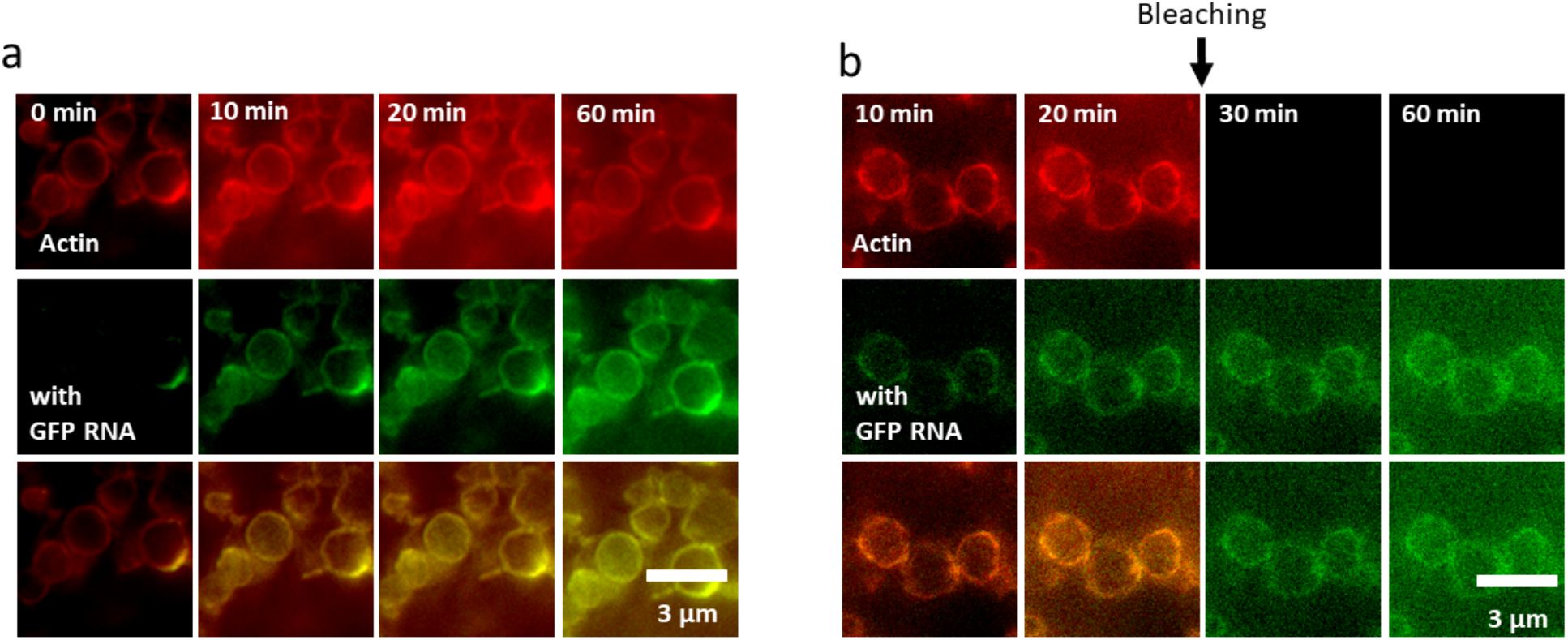
Actinosomes as efficient bioreactors. (**a**) Expression of GFP inside actinosomes by encapsulation of GFP-encoding mRNA and a cell-free *in vitro* translation machinery. As can be seen, GFP fluorescence increases over the course of an hour inside actinosomes while the background remains dark, indicating the protein expression is carried predominantly inside the containers. (**b**) To further clarify that the fluorescent signal is indeed coming from the GFP that is being produced, we bleached the ATTO-532 labelled actin fluorescence at 25 min, using high LED intensity (50%) for 5 s. This completely eliminated the signal from actin (the top row) but had no effect on the GFP fluorescence, which showed the same intensity as before and further increased over time. (**b**) Negative control (no GFP-encoding RNA but translation machinery is still present) showing no increase in fluorescence over the same duration.

## Discussion

In this paper, we have presented actinosomes: three-dimensional, cell-sized confinements with a boundary made up of polylysine polymers and actin filaments (Fig. 5). The unstructured proteinaceous shell provides a stable and porous boundary, allowing biochemical reactions to take place inside the container. Actinosomes are quick and easy to make, especially compared to the other containers such as liposomes and proteinosomes that are currently used to form synthetic cells. Furthermore, the use of initial condensate templates makes encapsulation of biomolecules particularly easy due to their intrinsic ability to concentrate a wide variety of biomolecules. The problem of poor encapsulation commonly encountered in bulk techniques is thus solved by using condensates as the starting material.

**Figure 5.**
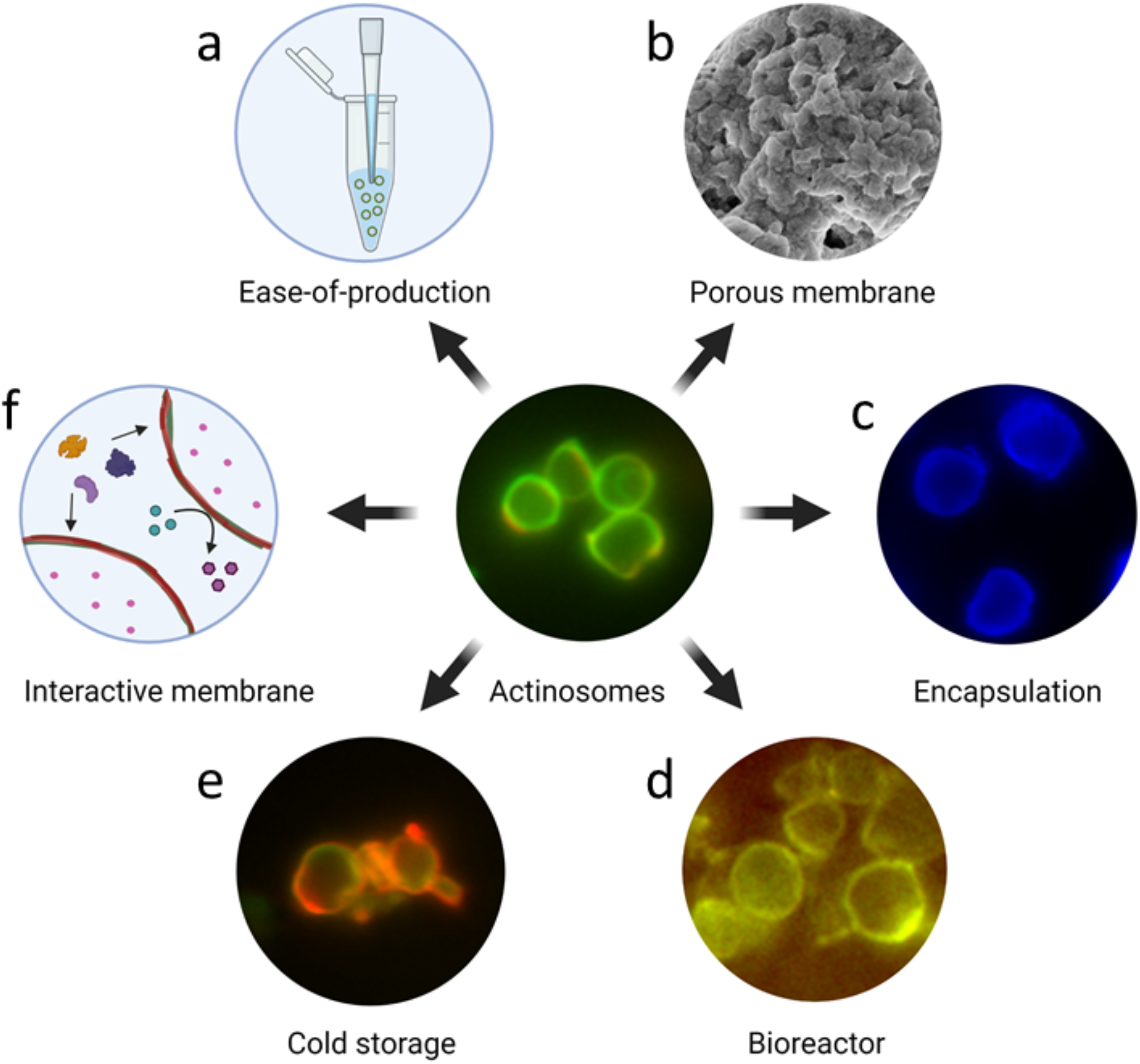
Salient feature of actinosomes. Actinosomes are synthetic confinements with a boundary made of polyK and actin filaments. Several properties make them potentially useful containers for synthetic cell research: (**a**) They are easy to produce without the need of any sophisticated set ups. (**b**) They have a porous surface (~0.5 nm sized openings) allowing molecules to pass through. (**c**) They can efficiently encapsulate biomolecules owing to the inherent sequestration capacity of condensates. (**d**) They have the capacity to act as a bioreactor to conduct complex reactions like protein translation. (**e**) They are stable and can be flash-frozen, stored at −80°C, and revived. (**f**) The actin-based boundary opens up the possibility to have an interactive membrane for recruiting other proteins, design signaling cascades, and form multicellular assemblies.

The current extensive use of microfluidic techniques to generate highly monodispersed containers and achieve efficient encapsulation also adds a significant amount of complexity to the production techniques^[2,49,50]^. An easy and robust process to produce micro-confinements can thus be useful for specific purposes and when such sophisticated set ups are not available. We have shown that condensate-templated actinosomes are straightforward to produce and are capable of carrying out complex biochemical reactions. While they might not be suitable for some specific features like growth and division, they are certainly appealing to be used as chemical nano factories and for studying the effect of confinement on biochemical processes. With regards to monodispersed samples, actinosomes were found to be surprisingly uniform in size (average diameter 3.1 ± 0.7 μm). We think this is due to nucleation, and subsequently coacervation, taking place homogenously throughout the solution and the droplets getting immediately stabilized by actin, further preventing their coalescence leading to a fairly homogenous size distribution.

We propose the following mechanism for actinosome formation: We begin with an initial homogenous solution of actin, polylysine, and other biomolecules that one wishes to sequester inside actinosomes. Upon addition of NTP mixture (ATP + GTP) to the solution, complex coacervation is induced between polylysine and NTP, forming coacervate droplets. Actin preferentially decorates the surface of the condensates, aided partially by the electrostatic interactions between net negatively charged actin protein and positively charged condensates and the viscous nature of the coacervate due to absence of salt. The other biomolecules (ones that are to be encapsulated inside), thanks to the intrinsic sequestering properties of condensates, are likely to get partitioned inside, or alternatively at the surface of, the condensate. It is important to note that at this point, actin stays in monomeric form as there are no Mg^2+^ ions present in the system which are necessary for polymerization. Addition of a salt-containing buffer (Mg^2+^ and KCl) triggers rapid actin polymerization at the expense of the ATP that is highly concentrated (~ 50 mM) in the condensates. Conversion of ATP into ADP and Pi (inorganic phosphate) leads to dynamic changes in the coacervate composition. However, polylysine polymers cannot readily diffuse outside and remain entangled within actin filaments to form an unstructured shell at the interface. This process continues till majority of the polylysine is co-localized with the actin at the surface. This eventually leads to dissolution of the original condensate droplet to ultimately form a micro-container, comprised of an aqueous lumen surrounded by a proteinaceous shell.

We observe that the addition of a monovalent salt (KCl) plays an important role in actinosome formation. In the initial low salt-content buffer, the condensates are predominantly elastic and very stable, owing to absence of any screening interactions. Addition of KCl weakens the electrostatic attractions between the coacervate components, possibly facilitating ATP consumption by actin and the outward movement of polylysine molecules to form an entangled mesh with actin filaments. Furthermore, addition of KCl presents a sort of hyperosmotic shock to the forming actinosomes with a semi-permeable interface. This seems to result in a water flux out of the actinosomes, possibly aiding to the distribution of the coacervate components at the interface. This logic is consistent with the different scenarios we observe as we change *R*: in case of low-enough GTP content (*R* ≤ 0.6), the hyperosmotic shock (Δ*c* ~ 200 mosm; Δ*P* = Δ*cRT* ~ 0.5 MPa) is too strong resulting in significant loss of the water content from the condensate, eventually resulting in a crumpled structure without any aqueous interior. At intermediate GTP content (*R* = 0.7 – 0.8), while there is efflux of water, the NTP concentration (~50 mM) is enough to sustain the osmotic pressure difference until the salt equilibrates. At high GTP content (R ≥ 0.9), there are no significant morphological changes as the actin does not polymerize readily due to lack of enough ATP and thus condensate components do not really change. Thus, actin polymerization at the expense of ATP inside the condensates and a sudden addition of salt together drives actinosome formation.

In conclusion, actinosomes are a novel addition as synthetic cell containers with many useful properties. They are easy-to-produce, require only basic lab equipment and commercially available relatively inexpensive proteins (Fig. 5a). They have a porous membrane with relatively large openings (~0.5 nm) allowing easy transport of molecules (Fig. 5b). Nonetheless, they can efficiently encapsulate biomolecules, especially charged ones like RNA (Fig. 5c). They can further carry out biochemical reactions inside (Fig. 5d). They can even be flash-frozen, stored at – 80 °C, and revived without losing structural integrity (Fig. 5e). Lastly, actin-based membrane presents interesting opportunities to functionalize these containers (Fig. 5f). For example, actin can interact with numerous actin-binding proteins to initiate specific reactions at the interface. This can be used in forming communicative networks within a population or even physically connecting the containers to form synthetic tissues. Such functionalities together with their highly biocompatible nature may allow actinosomes to interact with living cells and form hybrid interfaces. Further systematic research in these directions will reveal the true potential of these proteinaceous confinements and their use as scaffolds for synthetic cells.

## Materials and Methods

### Chemicals and proteins

Unlabeled poly-L-lysine (molecular weight 15-30 kDa) and fluorescently labelled FITC-poly-L-lysine (molecular weight 15-30 kDa) was purchased from Sigma-Aldrich. Individual nucleotides (ATP and GTP) were purchased from Thermo Scientific. Actin (rabbit skeletal muscle alpha actin) and fluorescent labelled ATT0 532-actin (rabbit skeletal muscle alpha actin) were purchased from Hypermol in the form of lyophilized powders. The composition of the reconstitution buffer to dissolve actin monomers was 2 mM Tris (pH 8.0), 0.4 mM ATP, 0.1 mM CaCl_2_ and 0.01 mM dithiothereitol. The end composition of the actin polymerization buffer was 0.01 M imidazole pH 7.4, 0.1 M KCl, and 2 mM MgCl2.

### Actinosome synthesis

The process of making actinosomes can be summed up in three distinct steps. 1) preparing actin polylysine mixture, 2) forming coacervates with coated actin 3) actin polymerization and coacervate dissolution. Step 1: Monomeric actin and polylysine were reconstituted in the actin reconstitution buffer, with final concentrations of 3 μM and 5.05 mg/ml, respectively. The pH 8 of the buffer is crucial for monomeric actin stability. Additionally, it keeps the polylysine polymers positively charged. For microscopic visualization, the sample was doped with 10% fluorescent labelled actin (0.3 μM) and 1% FITC-poly-L-lysine (0.05 mg/mL). Step 2: To trigger coacervation, 5 mM of NTP mixture (for example, 1.25 mM GTP and 3.75 mM ATP) was added to the solution and gently pipetted to mix thoroughly. Step 3: To make actinosomes, actin polymerization buffer was added to actin-coated coacervate solution. The sample was vortexed briefly to ensure sufficient mixing, followed by a short spin (1000 rpm for 5-10 seconds) to removes any aggregates. The last step significantly increased the actinosome yield.

### Zeta potential measurements

The net surface charge of the coacervate was determined by measuring the zeta potential at 25°C using the Malvern Zetasizer Nano instrument. The sample was diluted 1:20 and gently mixed prior to measurements. The zeta potential for each sample was determined by taking the average of five readings.

### ATP concentration measurements

To determine the NTP concentration required to obtain maximum amount of the condensate phase for a given polylysine concentration, we prepared buffered solutions (2 mM Tris (pH 7.4), 100 mM KCl, and 2 mM MgCl2) containing different concentration of ATP (from 1.25 mM to 25 mM) while keeping the polylysine concentration constant at 5 mg/ml. The solution was incubated at room temperature for 15 min to equilibrate. The condensed phase was separated from the dilute phase by centrifugation at 10,000 rpm for 5 minutes. The concentration of the free ATP in the dilute phase was evaluated by measuring its absorbance at 259 nm using the molar extinction coefficient of ATP (15,400 M^-1^ cm^-1^) using UV-Vis absorption spectroscopy (NanoDrop 2000/2000c Spectrophotometer, Thermo Scientific). The concentration of ATP inside the coacervates was calculated as *c_dense_* = (*c* − *c_dilute_f*)/(1 − *f*), where *c_dense_* and *c_dilute_* is the ATP concentration in dense and dilute phase respectively, and *f* is the volume fraction of the dilute phase. Concentration in the dilute phase, *c_dilute_*, was measured by absorbance as stated above. The fraction of the dilute phase, *f*, was estimated to be 0.9 by carefully removing the supernatant after centrifugation without disturbing the dense phase. For example, from a 40 μl sample, we estimated 36 μl to be the dilute phase.

### SEM microscopy

Surface of actinosomes was analyzed by scanning electron microscopy. Actinosomes were prepared and vacuum-dried at room temperature on electrically conductive carbon adhesive discs mounted on a metal stub. The dried sample were sputter-coated with Tungsten (to obtain a thin film of ~12 nm). The acquired images were taken at 10,000x magnification at 2 kV accelerating voltage and 13pA current (Fig. 3a, 3c, 3d) and at 35,000 x magnification at 5 kV accelerating voltage and 13 pA current (Fig. 3b).

### RNA expression in actinosomes

Capped and tailed messenger RNA (mRNA) template, encoding green fluorescent protein (GFP), was synthesized using the HiScribe T7 ARCA mRNA kit (New England Biolabs, Ipswich, MA) and a linearized double-stranded DNA (Supplementary Figure 8). The synthesized mRNA was purified using Monarch RNA Cleanup Kit (New England Biolabs, Ipswich, MA), thereby removing the template DNA. To ensure efficient encapsulation, GFP-mRNA (final concentration 100 ng/μl) along with *in vitro* translation machinery (Retic Lysate IVT™ Kit, Invitrogen,) was added along with actin and polylysine in step 1, prior to the addition of NTPs. This strategy allows efficient encapsulation of GFP-mRNA and translation machinery inside the actinosomes. Real-time expression of GFP was monitored by incubating actinosomes at 29°C using Okolab heating stage.

### Image analysis

Since the morphology of actinosomes are close to sphere, the size of the actinosomes were determined by fitting an ellipse using the Fitting Elipse function in Fiji. The obtained major and minor axes were used to determine the aspect ratio. For calculating the partition coefficient, the mean fluorescent intensity of actin and polylysine inside or at the surface of the coacervates (*I_dense_*) was measured for several coacervates, along with the mean fluorescent intensity outside the coacervates (*I_dnute_, n* = 10). The background intensity, *I_bg_*, was measured outside the sample. The corresponding partition coefficient was then calculated as 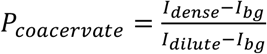.

## Acknowledgements

We thank Jasper van der Gucht for fruitful discussions. We thank Riccardo Antonelli and Thomas Kodger for help with scanning electron microscopy and Selena Koene for mRNA synthesis. The GFP-DNA template was a kind gift from Maria Forlenza and Mark Goldman (Aquaculture, Fisheries, and Immunology lab at Wageningen University). S.D. acknowledges financial support by the Innovation Program Microbiology grant (IPM-3) and by a ENW-KLEIN grant (OCENW.KLEIN.465) from the Dutch Research Council (NWO).

## Author Contributions

K.A.G. and S.D. conceived the idea, K.A.G. and L.L. performed the experiments and analysed the data. K.A.G. and S.D. wrote the paper.

