## Supplementary Information for "Actinosomes: condensate-templated proteinaceous containers for engineering synthetic cells"

**Supplementary figures**

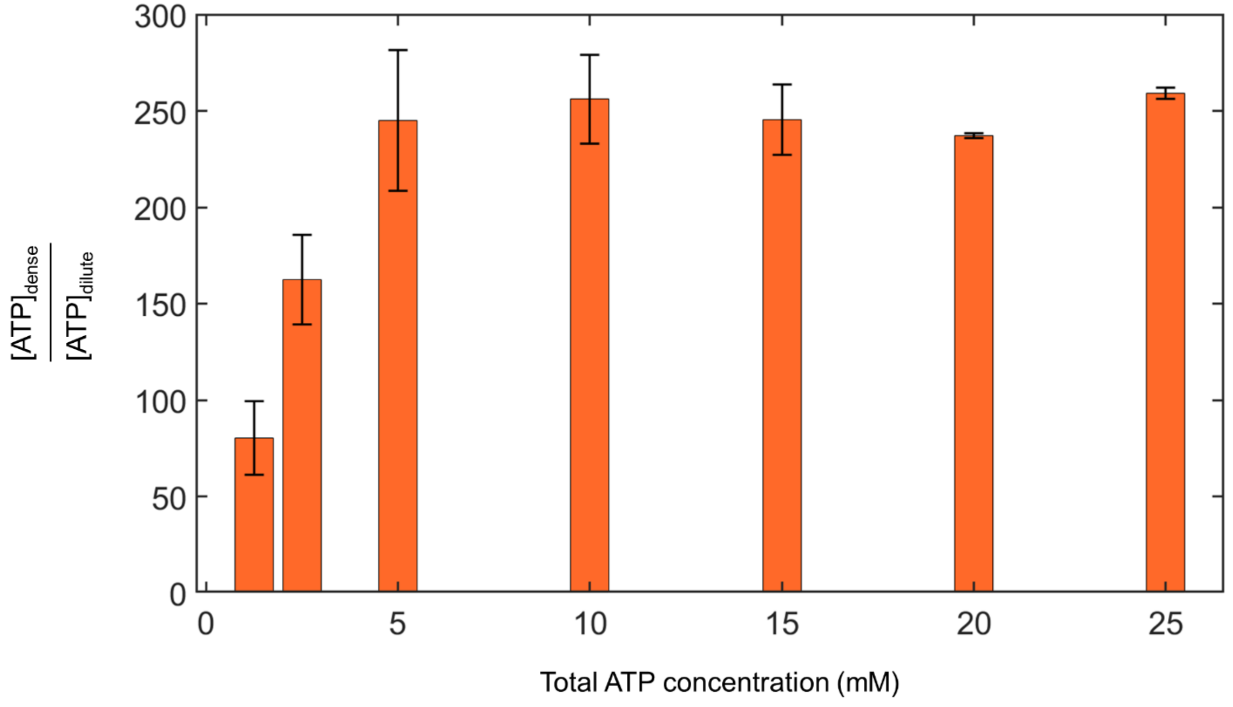


**Supplementary Figure 1. Determining the NTP concentration that leads to optimal coacervation with polyK.** Amount of ATP partitioned inside the dense phase at constant polylysine concentration of 5 mg/ml. The maximum partitioning (around 250-old) was observed for ATP concentration above 5.0 mM ($n$ = 3 experimental repeats, error bars indicate standard deviation; see Methods for details).

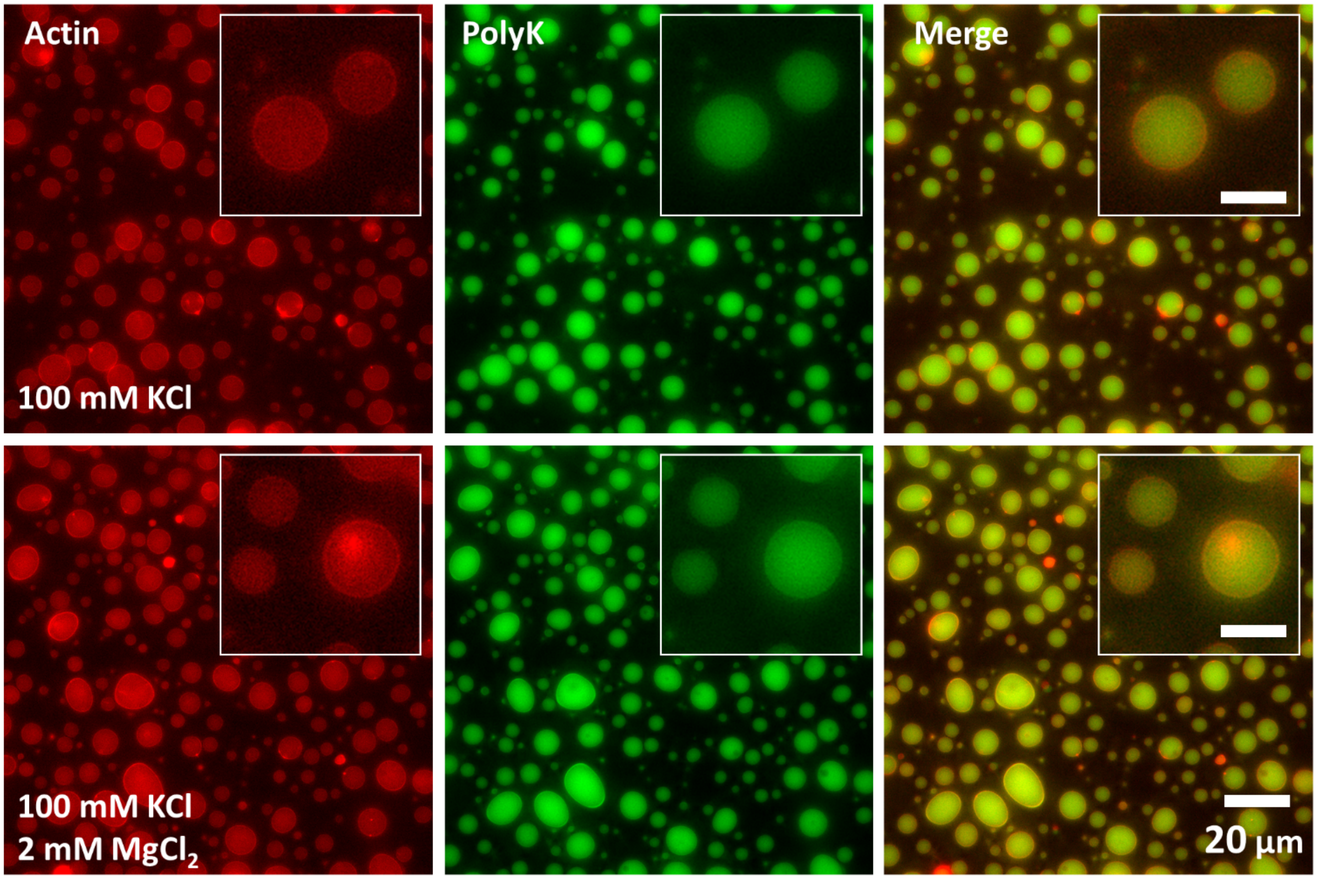


**Supplementary Figure 2. Presence of KCl in the initial reaction mixture leads to homogenous partitioning of actin in the coacervates and inhibits actinosome formation.** Coacervate droplets composed of polyK/NTP ($R$ = 0.7) made in presence of KCl sequester the actin within the coacervates (top), as opposed to its localization at the interface in absence of KCl (Fig. 1). Triggering actin polymerization by adding $MgCl_2$ does not result in actinosome formation (bottom). Scale bars in the insets represent 3 µm.

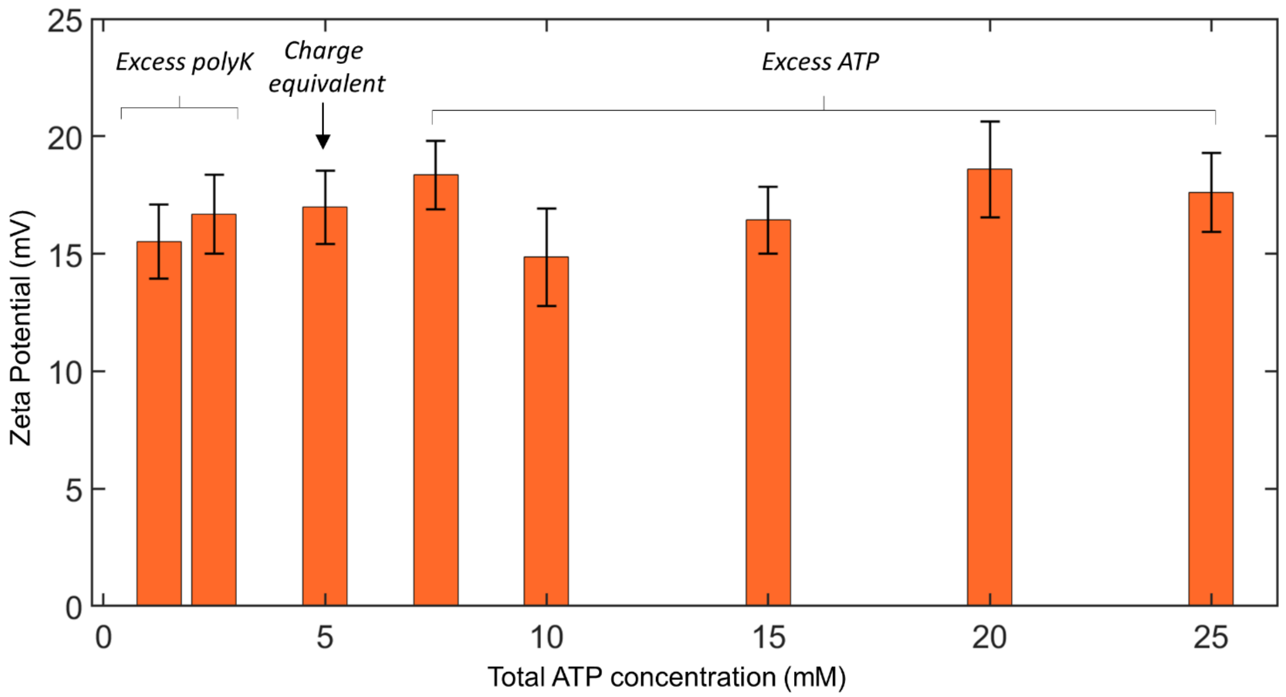


**Supplementary Figure 3. Surface charge of polyK/ATP coacervates stays constant over different ATP:polyK ratios.** Zeta potential of the polyK/ATP coacervate droplets was measured over a concentration range of 1.25 – 25 mM, with polyK concentration kept constant at 5 mg/mL. The measurements covered three regimes: excess polyK (positively charged polymer), charge equivalent state (net neutral), and excess ATP (negatively charged multivalent molecule). The values stayed relatively constant (15.0 ± 1.6 mV) over the entire ATP concentration range, suggesting net positive surface charge on the coacervate droplets irrespective of limiting or excess ATP concentrations ($n$ = 3 experimental repeats with each measurement being an average of 5 individual runs. Error bars indicate standard deviation.).

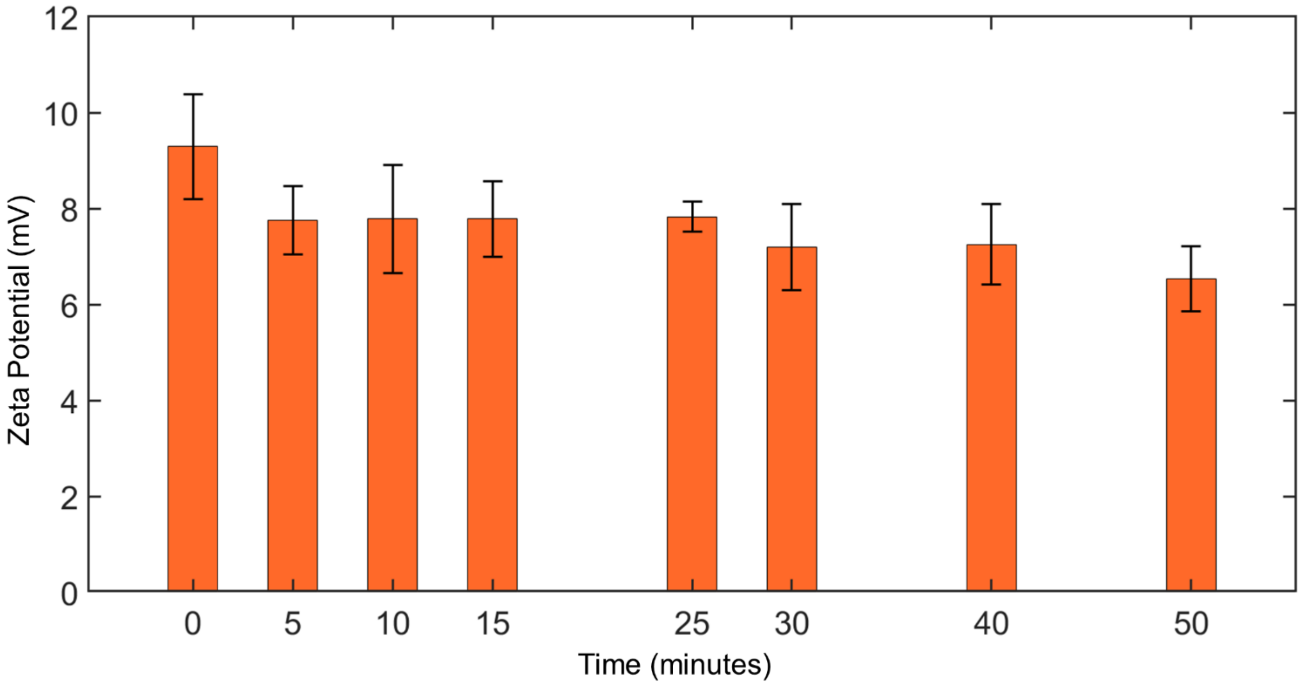


**Supplementary Figure 4. Surface charge of polyK/ATP coacervates in presence of actin monomers over time.** Zeta potential of polyK/ATP coacervate droplets was measured at different time points after the addition of actin. PolyK, ATP, and actin concentrations were respectively kept at 5 mg/mL, 5.4 mM, and 3 µM respectively; polymerization conditions were used. Immediately after the addition of actin, the surface charge was clearly lowered (< 10 mV) compared to when actin was absent (15.0 ± 1.6 mV; Supplementary Figure 2). This indicates efficient accumulation of actin at the surface. The surface charge was further decreased to 7.8 ± 0.7 mV after 5 minutes and then remailed fairly constant over the entire time duration (*n* = 3 experimental repeats with each measurement being an average of 5 individual runs. Error bars indicate standard deviation.).

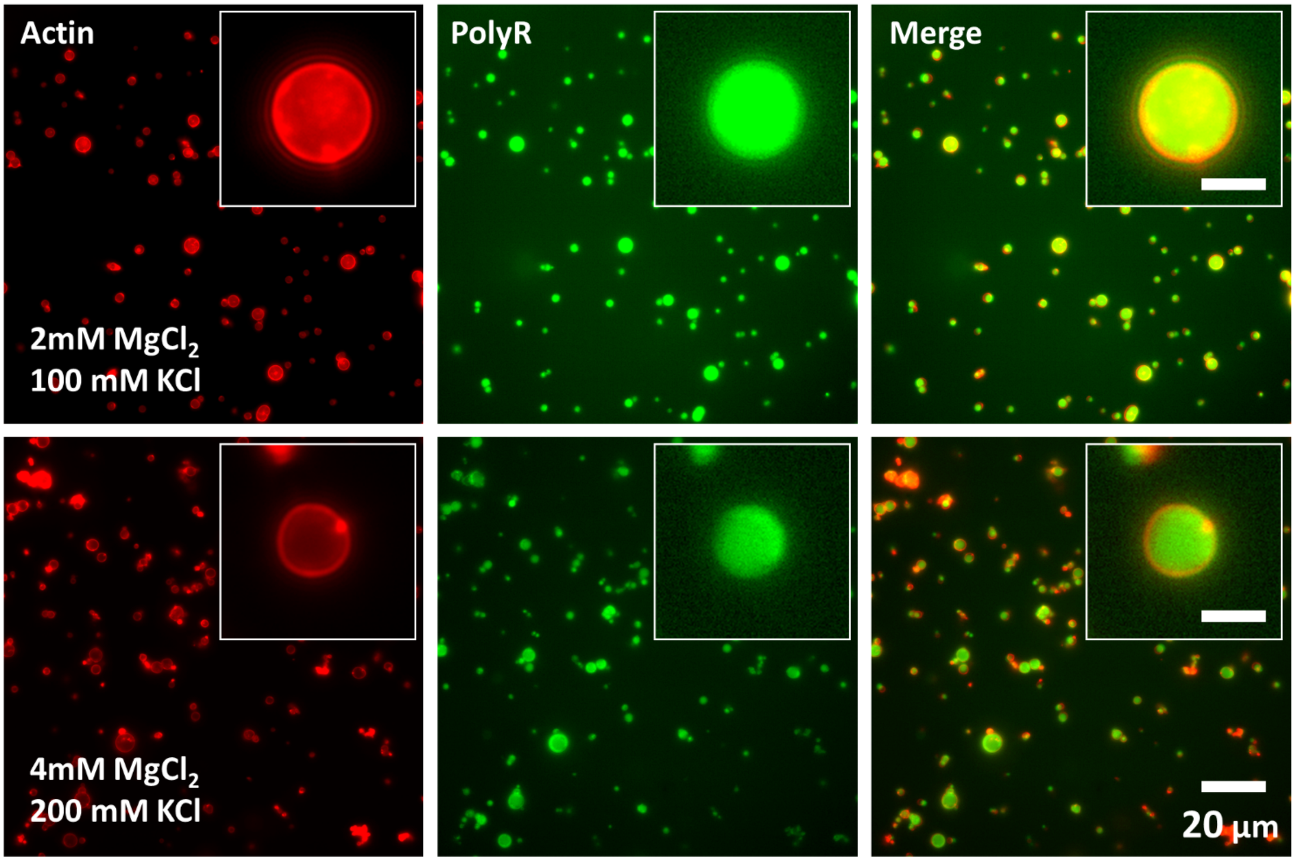


**Supplementary Figure 5. PolyR/NTP coacervates do not form actinosomes.** Coacervate droplets composed of polyR/NTP (*R* = 0.7) do not form actinosomes at similar buffer conditions (top) and even after doubling the concentrations of $Mg^{2+}$ and KCl (bottom). Scale bars in the insets represent 3 µm.

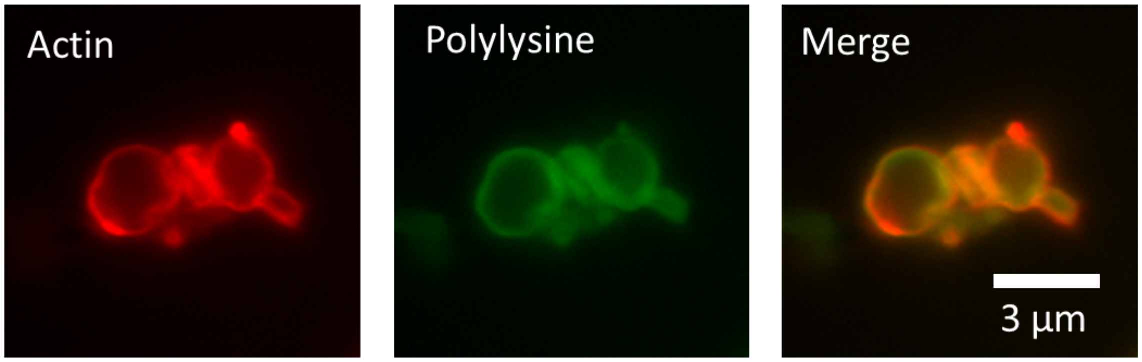


**Supplementary Figure 6. Actinosomes can be frozen and revived.** Freshly prepared actinosomes were flash-frozen using liquid nitrogen and stored at −80°C (for 24 hours) and subsequently restored at room temperature. Fluorescence images confirming the structural integrity of actinosomes.

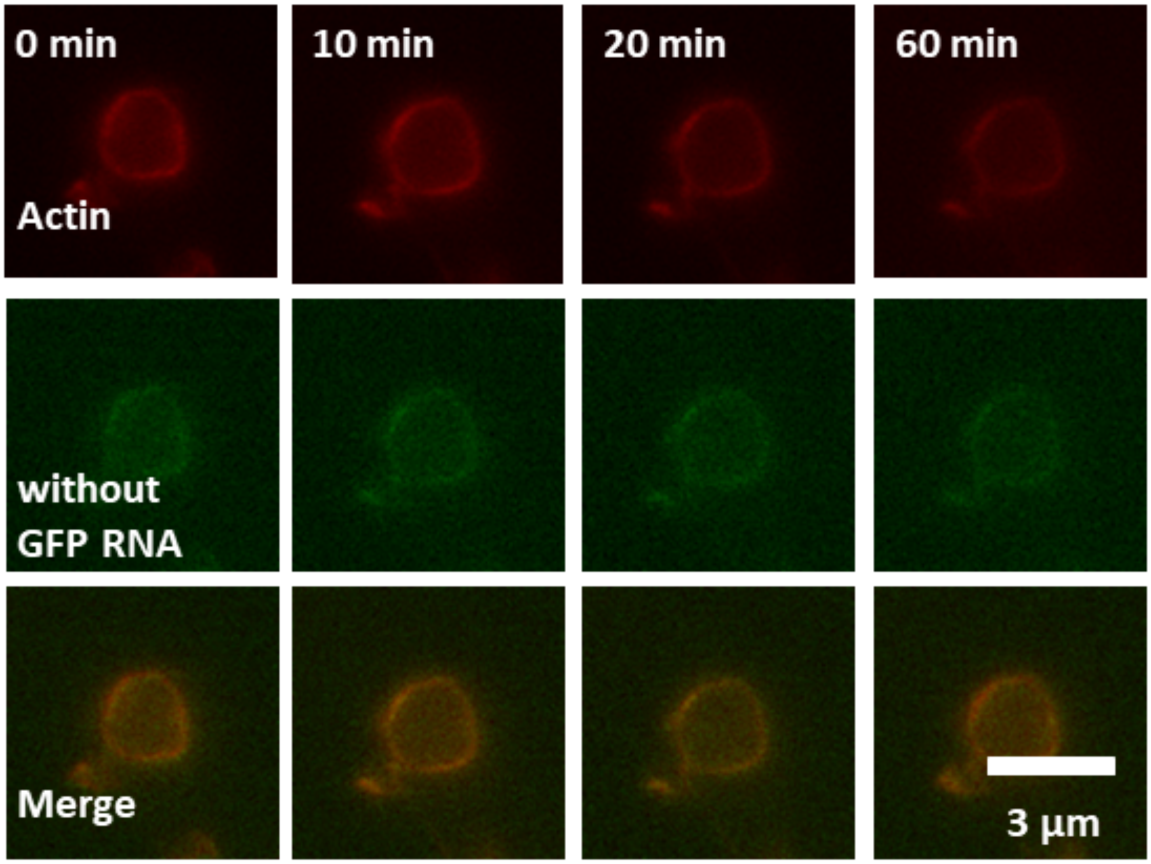


**Supplementary Figure 7. Actinosome encapsulating protein expression machinery in absence of GFP-RNA.** Negative control (no GFP-encoding RNA but translation machinery is still present) showing no increase in the GFP fluorescence intensity over the time duration of an hour.

T7 Promotor

CCACTGCTTACTGGCTTATCGAAATTAATACGACTCACTATAGGGAGACCCAAGCTGGCTAGCGTTTAAACTTAAGCTTGGTACCGAGCTCGGATCCACTAGTCCAGTGTGGTGGAATTCA

GFP encoding

ACATGGTGAGCAAGGGCGAGGAGCTGTTCACCGGGGTGGTGCCCATCCTGGTCGAGCTGGACGGCGACGTAAACGGCCACAAGTTCAGCGTGTCCGGCGAGGGCGAGGGCGATGCC

ACCTACGGCAAGCTGACCCTGAAGTTCATCTGCACCACCGGCAAGCTGCCCGTGCCCTGGCCCACCCTCGTGACCACCCTGACCTACGGCGTGCAGTGCTTCAGCCGCTACCCCGACCAC

ATGAAGCAGCACGACTTCTTCAAGTCCGCCATGCCCGAAGGCTACGTCCAGGAGCGCACCATCTTCTTCAAGGACGACGGCAACTACAAGACCCGCGCCGAGGTGAAGTTCGAGGGCG

ACACCCTGGTGAACCGCATCGAGCTGAAGGGCATCGACTTCAAGGAGGACGGCAACATCCTGGGGCACAAGCTGGAGTACAACTACAACAGCCACAACGTCTATATCATGGCCGACAA

GCAGAAGAACGGCATCAAGGTGAACTTCAAGATCCGCCACAACATCGAGGACGGCAGCGTGCAGCTCGCCGACCACTACCAGCAGAACACCCCCATCGGCGACGGCCCCGTGCTGCTG

CCCGACAACCACTACCTGAGCACCCAGTCCGCCCTGAGCAAAGACCCCAACGAGAAGCGCGATCACATGGTCCTGCTGGAGTTCGTGACCGCCGCCGGGATCACTCTCGGCATGGACG

AGCTGTACAAGTAAGTCTAGAGGGCCCGTTTAAACCCGCTGATCAGCCTCGACTGTGCCTTCTAGTTGCCAGCCATCTGTTGTTTGCCCCTCCCCCGTGCCTTCCTTGACCCTGGAAGGTG

CCACTCCCACTGTCCTTTCCTAATAAAATGAGGAAATTGCATCGCATTGTCTGAGTAGGTGTCATTCT

**Supplementary Figure 8. DNA template used for GFP production.** Template DNA encoding green fluorescent protein (highlighted in green) under the bacterial T7 promotor (highlighted cyan) that was used for cell-free *in vitro* synthesis of mRNA.
